# Genetic analysis of *C. elegans* Haspin-like genes shows that *hasp-1* plays multiple roles in the germline

**DOI:** 10.1101/2022.02.12.480216

**Authors:** Jommel Macaraeg, Isaac Reinhard, Matthew Ward, Danielle Carmeci, Madison Stanaway, Amy Moore, Ethan Hagmann, Katherine Brown, David J Wynne

## Abstract

Haspin is a histone kinase that promotes error-free chromosome segregation by recruiting the Chromosomal Passenger Complex (CPC) to mitotic and meiotic chromosomes. Haspin remains less well studied than other M-phase kinases and the models explaining Haspin function have been developed primarily in mitotic cells. Here, we generate new mutations in the *C. elegans* Haspin homologs *hasp-1* and *hasp-2* and characterize their phenotypes. We show that *hasp-1* is responsible for all predicted functions of Haspin and that loss of function of *hasp-1* using classical and conditional alleles produces defects in germline stem cell proliferation, spermatogenesis, and confirms its role in oocyte meiosis. Genetic analysis suggests *hasp-1* acts downstream of the Polo-like kinase *plk-2* and shows synthetic interactions between *hasp-1* and two genes expected to promote recruitment of the CPC by a parallel pathway that depends on the kinase Bub1. This work adds to the growing understanding of Haspin function by characterizing a variety of roles in an intact animal.

**Summary statement:** We characterize new mutations in the *C. elegans* homologs of the histone kinase Haspin and show roles in spermatogenesis, germline proliferation and genetic interactions during oocyte meiosis.

## Introduction

Haspin is a protein kinase that is important for the correct segregation of chromosomes in mitosis and meiosis (Higgins, 2010;Hindriksen et al., 2017). It is one of many kinases that coordinate the events of M phase, such as Cyclin-dependent kinase 1 (Cdk1), Polo-like kinase 1 (Plk1), and Aurora B. Intricate regulatory interactions have been identified among these proteins so a complete understanding of this network requires specific manipulations and benefits from being studied using diverse model systems. Haspin inhibitors hold the potential of being effective chemotherapy drugs because features of Haspin’s kinase domain are unique among eukaryotic protein kinases (Dominguez-Brauer et al., 2015;Eswaran et al., 2009;Villa et al., 2009). Haspin phosphorylates Histone H3 on threonine 3 (H3T3), which is generally thought to be the single most important substrate for Haspin function during mitosis and meiosis (Dai et al., 2005). The possibility of a single critical target has made Haspin inhibitors attractive chemotherapeutics because of the potential for fewer off target effects (Amoussou et al., 2018). However, there is some evidence that Haspin may have other targets, including from proteomic studies with Haspin inhibitor treatments, which suggested the histone variant macroH2A and the centromeric protein CENP-C may be direct targets (Maiolica et al., 2014). The testis-specific histone H2A protein TH2A has been shown to be phosphorylated by Haspin *in vivo* and the cohesion regulator WAPL is phosphorylated by Haspin *in vitro* (Hada et al., 2017;Liang et al., 2018). The challenge of determining the direct targets of a kinase and the discrepancies that can occur between *in vitro* and *in vivo* kinase activities warrant additional analysis of Haspin function using diverse *in vivo* systems.

Haspin has two overlapping functions in mitosis at the inner centromere, a distinct region of mitotic chromosomes in between the sister centromeres. Haspin helps protect chromosome cohesion by binding to the cohesion-associated protein Pds5 and inhibiting the cohesin-removing protein WAPL (Dai et al., 2006;Goto et al., 2017;Liang et al., 2018;Yamagishi et al., 2010;Zhou et al., 2017). Haspin-dependent phosphorylation of H3T3 (H3T3ph) directly recruits the Chromosomal Passenger Complex (CPC), which plays a variety of roles that promote error-free chromosome segregation (Carmena et al., 2012). The CPC is composed of four subunits, the Aurora B kinase and three accessory subunits INCNEP, Borealin/Dasra, and Survivin, which directly binds H3T3ph (Kelly et al., 2010; Wang, F. et al., 2010; Yamagishi et al., 2010). These two mechanisms help explain the phenotypes seen for Haspin knockdown in mitotic cells, which include misaligned chromosomes, loss of cohesion, and spindle defects (Dai and Higgins, 2005;Dai et al., 2006;Dai et al., 2009), but the effects of Haspin deletion in unperturbed mitotic cells are relatively mild (Hadders et al., 2020). Haspin is activated by phosphorylation of multiple sites in its long, disordered amino-terminal domain, which has been shown to be a target of Plk1, Cdk1, Aurora B, and autophosphorylation (Ghenoiu, C. et al., 2013; Wang, F. et al., 2011; Zhou et al., 2014). It remains to be seen how important these regulatory mechanisms are in a variety of cell types and whether they are conserved in more distantly related species.

The relatively mild phenotype of Haspin loss in mitosis is due to partially redundant function with the Spindle Assembly Checkpoint kinase Bub1, which promotes CPC localization through a second kinetochore-associated mechanism (Yamagishi et al., 2010). Bub1-dependent phosphorylation of histone H2A recruits shugoshin proteins, which recruit the Borealin/Dasra subunit of the CPC (Tsukahara et al., 2010). There has been uncertainty about the interplay between these two parallel pathways (Hindriksen et al., 2017), but recent work in human tissue culture cells supported the model that both pathways can independently recruit the CPC to chromatin (Hadders et al., 2020). The functional significance of different CPC recruitment mechanisms and therefore the existence of distinguishable pools of CPC is still being investigated. Moreover, work is ongoing to understand the interplay between kinetochore-dependent signaling molecules and cohesion regulators. There is a need for better understanding of the extent to which CPC recruitment and cohesion protection mechanisms are altered in different cell types and in different organisms. Cancer cells have been shown to lose the ability to recruit the CPC preferentially to misaligned chromosomes, so understanding how CPC recruitment mechanisms are altered is important for future cancer treatments (Salimian et al., 2011).

In contrast to mitosis, relatively little is known about Haspin function in meiosis and somewhat contradictory findings have been reported in different organisms. A Haspin knockout mouse is fertile and shows no obvious developmental phenotypes other than some disordered cells in the testis (Shimada et al., 2016). In contrast, Haspin inhibitors cause defects in spindle pole clustering and failure to complete meiosis I in mouse spermatocytes, as well as incorrect kinetochore-microtubule attachments that lead to aneuploidy (Balboula et al., 2016;Nguyen et al., 2014). Surprisingly, Haspin inhibition also caused a loss of condensin on meiotic chromosomes but no defects in cohesion were seen (Nguyen et al., 2014). The meiotic bivalent is a cruciform structure made up of 4 chromosomes, and Haspin inhibition clearly disrupted CPC localization on the axes between chromosomes but left kinetochore-proximal pools untouched (Nguyen et al., 2014). This phenotype is very similar to the kinetochore CPC pools seen after mitotic after Haspin loss, but over a ten-fold larger length scale (Bekier et al., 2015).

In *Drosophila*, Haspin has been shown to affect transcription during interphase and in histone segregation in germline stem cells (Fresan et al., 2020;Xie et al., 2015). However, loss of function of Haspin does not cause viability defects or meiotic nondisjunction in *Drosophila*, and Haspin-dependent recruitment of the CPC in meiosis is not as important as it is in mouse spermatocytes (Wang, L. I. et al., 2021). The discrepancies in Haspin’s role in meiosis among animals as well as its additional functions that can be more easily characterized in some systems justify further study of the protein’s function in model organisms.

*C. elegans* is a powerful system to study chromosome segregation in both mitosis and meiosis because there are many genetic and cytological tools available. Mutant alleles in cell cycle regulatory genes are readily available and easily combined by crosses. Individual hermaphrodites produce over 300 progeny, which makes fecundity measurements robust enough to distinguish small differences and cell division can be monitored through the transparent bodies and eggs of live animals. The *C. elegans* germline is organized so that nuclei move in one direction as they progress through mitotic stem cell divisions and then meiosis, with spermatogenesis and oogenesis occurring at specific life stages in hermaphrodites (Hubbard, E. J. A. Greenstein, D., 2005). Finally, the sex determination system, in which XO males are produced by meiotic nondisjunction in XX hermaphrodites, facilitates analysis of meiotic chromosome segregation and can distinguish meiotic defects from mitotic (Hodgkin et al., 1979).

Here, we set out to use *C. elegans* to better understand the function of Haspin-related genes in the context of a whole animal. To our knowledge, only a single previous report has tested the function of Haspin (hasp) proteins in this organism. In *C. elegans*, the mechanisms that regulate differential cohesion loss in meiosis have been rigorously studied. Unlike mouse oocytes where the CPC is found on both chromosome axes, the CPC is restricted to the “short arm” of the meiotic bivalent in oocytes, the axis in between the homologous chromosomes, where it is essential for anaphase I chromosome segregation (Kaitna et al., 2002;Rogers et al., 2002). The *C. elegans* Aurora B homolog AIR-2 phosphorylates the meiosis-specific cohesin REC-8, which leads to the selective loss of cohesion between homologs in meiosis I (Ferrandiz et al., 2018;Rogers et al., 2002). The mechanisms that restrict the CPC to this region in *C. elegans* have also been well studied. The nematode-specific protein LAB-1 plays a role analogous to that of shugoshin in other systems by protecting cohesion in meiosis I (de Carvalho et al., 2008). Like shugoshin proteins, LAB-1 recruits a phosphatase that antagonizes phosphorylation, including both H3T3ph and the AIR-2-dependent phosphorylation of H3 (Ferrandiz et al., 2018;Tzur et al., 2012). Recent work described a new model for CPC recruitment in *C. elegans* in which the spatial regulation of the CPC is primarily controlled by signaling from HASP-1 protein that is localized by the meiotic chromosome axis, while temporal regulation of the CPC depends on activity of the Cdk1 homolog, *cdk-1* (Ferrandiz et al., 2018). However, a number of questions about the function of Haspin-related genes in *C. elegans* remain. *C. elegans* has two genes similar to Haspin, *hasp-1* and *hasp-2* and it has not been tested whether *hasp-2* plays an important role. Previous studies in *C. elegans* focused on oocytes so it is not clear whether *hasp-1* is important for spermatogenesis. In addition, it is not known whether the Bub1-dependent pathway functions in parallel with *hasp-1* to recruit the CPC during meiosis, or if activation of *hasp-1* by Plk1 homologs is conserved in *C. elegans.*

In this study, we analyzed both classical and conditional mutations in the Haspin-related genes in *C. elegans, hasp-1* and *hasp-2.* We found that a deletion mutation in *hasp-1* caused sterility due to a defect in germline stem cell proliferation and we used a conditional *hasp-1* mutation to show that *hasp-1* acts in spermatogenesis as well as oogenesis. Depletion of *hasp-1* caused meiotic segregation errors and maternal embryonic lethality. Combining *hasp-1* depletion with other mutant backgrounds, we showed that *hasp-1* is likely to function downstream of the Plk1 homolog *plk-2*, which is consistent with it being activated by *plk-2.* In contrast, we found that mutations in Bub1 pathway genes caused strong synthetic interaction phenotypes when combined with *hasp-1* knockdown, suggesting that the parallel activity of the Bub1 pathway is required in *C. elegans* meiosis. Our results confirm that many of the regulatory mechanisms described for Haspin in other systems are conserved in *C. elegans* and that this model system will be useful to further test hypotheses for Haspin function and regulation in a variety of cell types in an intact animal.

## Results

### A deletion mutation in hasp-1 causes sterility due to a lack of germline stem cell proliferation

The Haspin gene has undergone an unusual expansion in the *Caenorhabditis* lineage with more than eight predicted proteins that have some homology to the vertebrate Haspin kinase domain (Higgins, 2001). Two of these paralogous genes have the highest homology to human Haspin and have been named *hasp-1* and *hasp-2.* A *hasp-1* deletion allele, *hasp-1(tm3858)* generated by the Japanese National Bioresourse Project for *C. elegans* (Gengyo-Ando and Mitani, 2000) had not yet been characterized to our knowledge. The *hasp-1(tm3848)* mutation is predicted to generate an in-frame deletion of 155 amino acids that include the beginning of the well-conserved kinase domain and a lysine residue (K644) required for kinase activity (**Fig. 1A**, (Villa et al., 2009). We verified the deletion breakpoints (data not shown) and outcrossed the mutation prior to analysis of the phenotype. Hermaphrodites homozygous for *hasp-1(tm3858)* grew to adulthood but were sterile with a slight protruding vulva (**Fig. 1B**). Sterility and vulval defects are common phenotypes for loss of function of genes involved in mitosis in *C. elegans* (O’Connell et al., 1998). The area around the vulva had a clear appearance due to the failure to produce embryos filling the uterus. We DAPI stained whole animals to visualize the gonad and found a dramatic lack of germline nuclei in adult animals (**Fig. 1C**). We assumed that the lack of germline nuclei was due to a failure of mitotic proliferation of germline stem cells, but we also considered the possibility of defects in somatic gonad development. To better characterize the germline defect, we crossed *hasp-1(tm3858)* into a background containing fluorescent fusion proteins marking the distal tip cell (DTC) and histone H2A, which allowed us to visualize the ends of the gonad arms and the nuclei inside (**Fig. 1D**). In synchronized animals 48-54 hours after egg laying, all DTCs in both mutant and control animals had successfully completed the first phase of migration by moving away from the vulva along the ventral side of the animal. However, by this stage it was clear that the germline stem cell proliferation that normally occurs during DTC migration had not occurred. In the mutants, fewer than ten nuclei were present and only in the distal gonad region while control gonads contained over a hundred nuclei. Nuclei in mutant gonads also varied in size with some much larger than in control animals (yellow arrowheads, **Fig. 1D**). In controls, some DTCs had also completed the later phases of migration by turning and moving back toward the vulva along the dorsal side, but in *hasp-1(tm3858)* animals, DTCs had at most completed only the first turn in migration in this time period. Because the germline proliferation defect was much more severe than the delay in DTC migration, these data support the hypothesis that a defect in proliferation is the cause of sterility in these animals. Because of the lack of germline in the *hasp-1(tm3858)* mutant, further study of *hasp-1* required a different approach to *hasp-1* knockdown.

**Figure 1.**
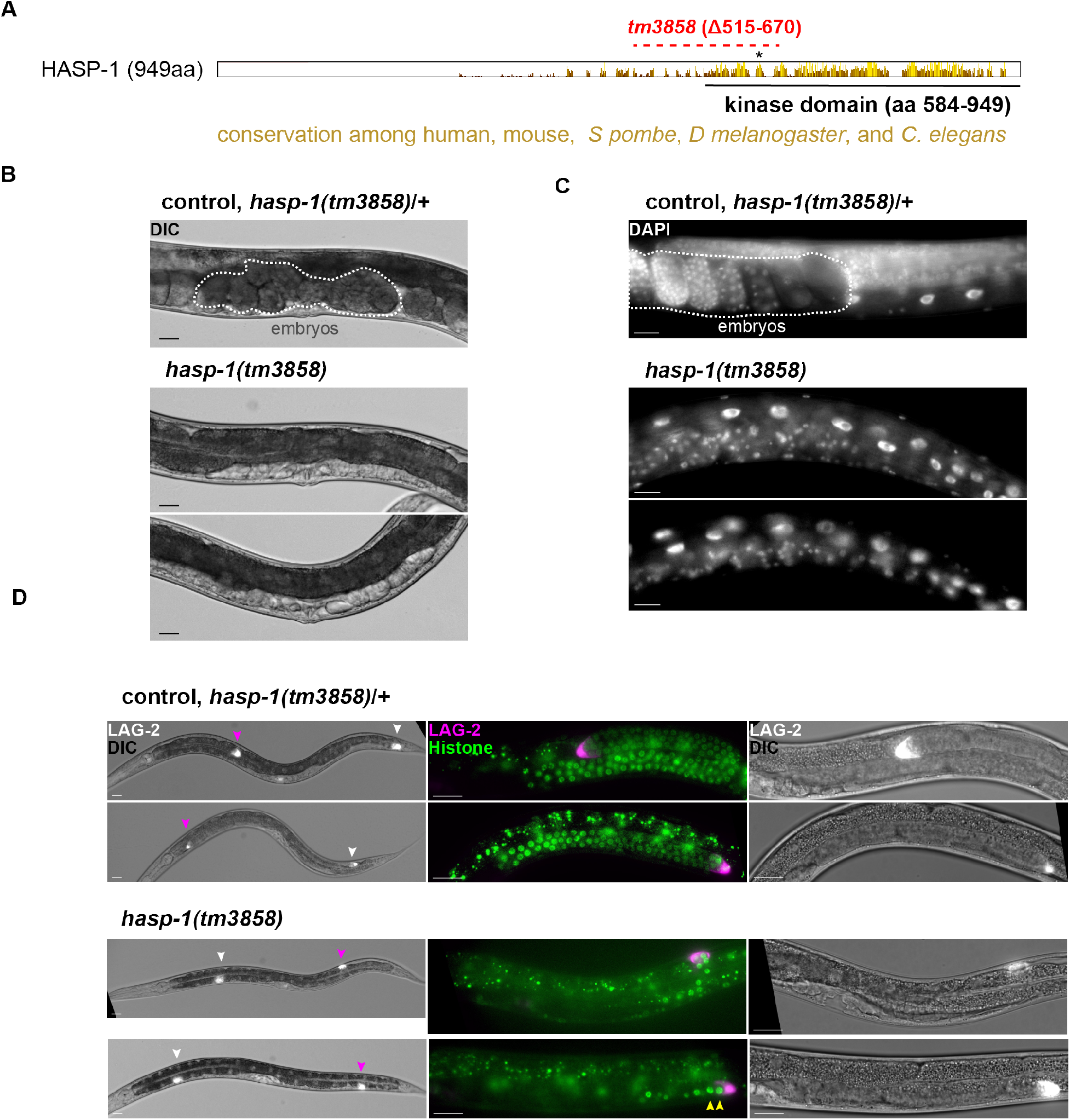
A putative null mutation in *hasp-1* causes sterility due to a failure of germline stem cell proliferation. A) schematic of HASP-1 protein showing locations of conservation among human, mouse, *S. pombe*, and *D. melanogaster* Haspin homologs. Lysing 644 (K644) is shown with an asterisk and the dotted red line indicates the in-frame deletion caused by the *hasp-1(tm3858)* allele. Morphology of *hasp-1(tm3858)* and control animals is shown using B) Nomarski, C) DAPI staining and D) fluorescent markers for the distal tip cell, which marks the end of the developing gonad arms (LAG-2::GFP, white and magenta arrowheads), and histones (green). Embryos are circled in white dotted lines in B-C to highlight the lack of embryos in the mutant. In D, the right two panels are higher magnification images of the gonad arm marked with the magenta arrowhead in the leftmost panel, oriented with the ventral side down and the vulva to the left. Yellow arrowheads show enlarged nuclei in a mutant gonad. Scale bars = 20 μm.

### A nonsense mutation in hasp-2 has no obvious phenotype

The *hasp-1* and *hasp-2* kinase domains have high identity (77%) and residues known to be required for kinase activity are conserved, including lysine and glutamate residues essential for ATPase activity as well as two aspartate residues essential for catalysis and magnesium binding (Manning et al., 2002;Villa et al., 2009). However, the *hasp-2* gene is predicted to produce a much smaller protein than *hasp-1* and to be made up almost entirely of the kinase domain (**Fig. 2A**). Thus, many of the regulatory mechanisms that have been characterized in the N-terminal portion of Haspin genes in other organisms are unlikely to be shared by *hasp-2.* For example, there are no Cdk1 phosphorylation sites (STP sequences) outside of the *hasp-2* kinase domain, which are expected to activate the kinase (Ghenoiu, Cristina et al., 2013) and no tyrosine residues, which are conserved in motifs required for Pds5 interaction (Goto et al., 2017). *hasp-1* has been suggested to have a divergent Pds5 interaction motif, but this has not yet been experimentally validated in *C. elegans* (Goto et al., 2017). Based on RNA-seq data available through Wormbase, *hasp-2* expression is highest in males and in L4 hermaphrodites, in contrast to *hasp-1*, which peaks in the early embryo. The expression data suggest the possibility that *hasp-2* may have evolved specificity for a role during spermatogenesis. We were interested in this possibility due to the high expression of Haspin in mouse testis relative to other proliferative tissues (Tanaka et al., 1999).

**Figure 2.**
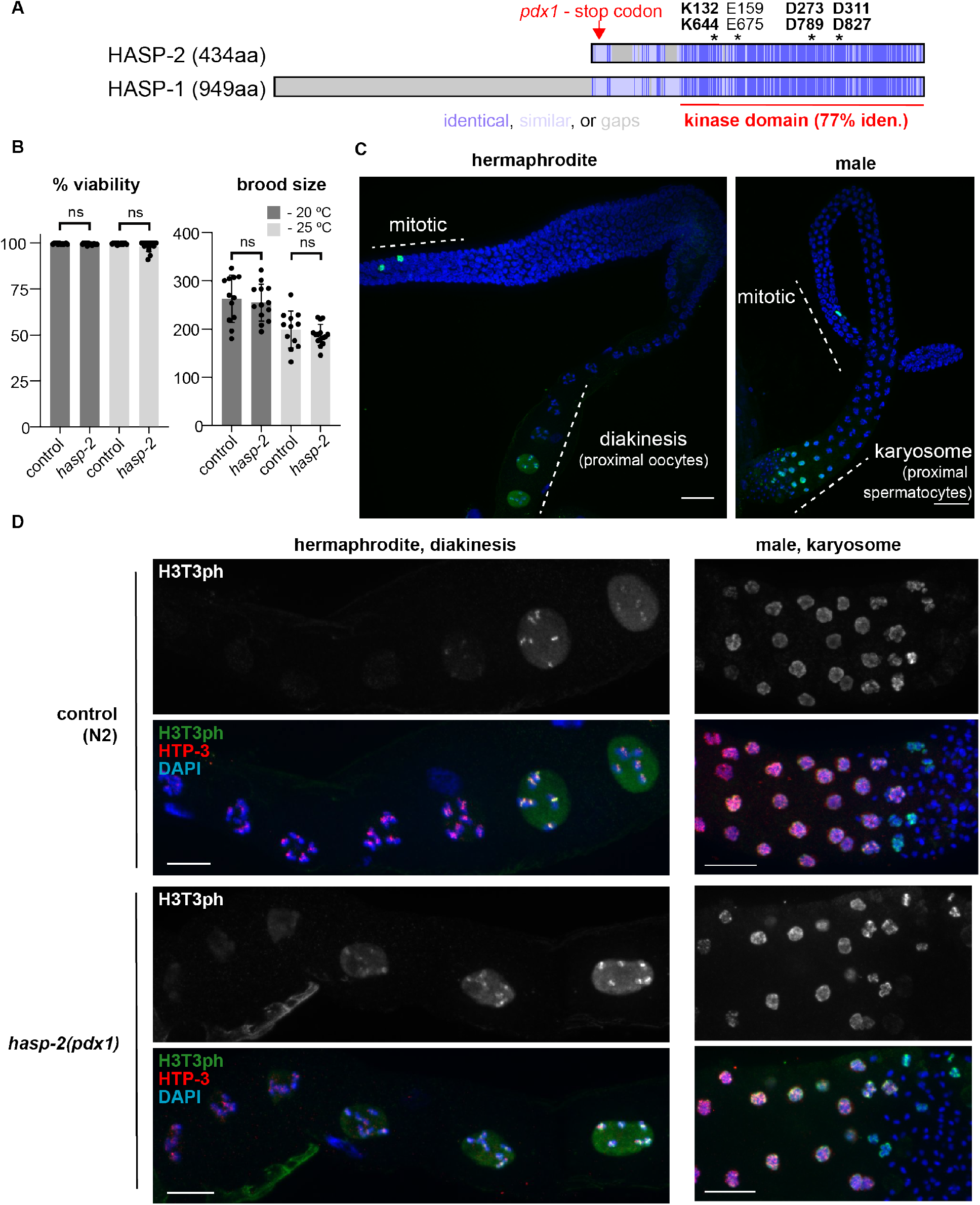
A *hasp-2* putative null mutation has no effect on brood or H3T3 phosphorylation. A) Schematic of HASP-1 and HASP-2 predicted proteins showing locations of conserved sequences (blue) and residues essential for kinase activity (asterisks). B) Average embryonic viability and brood size measurements for whole broods from control (N2) and *hasp-2(pdx1)* animals raised at 20 or 25°C (error bars show standard deviation). ns = p>0.05 by Mann Whitney test. C) H3T3ph staining in whole dissected gonads from a hermaphrodite (left) and male (right). Scale bar = 20 μm D) Higher magnification images as in C showing the nuclei in the proximal region of the gonad. Scale bar = 10 μm.

No previously existing mutation in *hasp-2* was available, so we used CRISPR/Cas9 to introduce a small insertion early in the first exon that generated a nonsense mutation followed by a frame shift (**Fig. 2A**). After verifying and outcrossing the allele, we found that animals homozygous for *hasp-2(pdx1)* had normal embryonic viability and brood sizes at both 20°C and 25°C (**Fig. 2B**) and no apparent morphological defects. We examined H3T3ph staining in the germline of *hasp2(pdx1)* mutants and saw robust signal in mitotic nuclei and in the proximal gonad regions in both hermaphrodites and males (**Fig. 2C,D**). Assuming that the *hasp-2(pdx1)* allele is acting as a null mutation, this and the results of double mutant strains discussed below suggest that *hasp-2* does not promote H3T3ph in the *C. elegans* germline. The severe phenotype caused by *hasp-1* loss of function (Fig. 1) and the lack of phenotypes associated with *hasp-2* mutation suggest that *hasp-2* is unlikely to substitute for *hasp-1* activity in *hasp-1* mutant strains and so we focused our attention on better characterizing the loss of *hasp-1.* We also note that our immunofluorescence experiments did not detect H3T3ph in any other nuclei in the adult hermaphrodite germline, such as mature sperm, suggesting that HASP-1 activity may be limited to meiosis in spermatocytes.

### Conditional degradation of HASP-1 causes maternal effect embryonic lethality without affecting germline proliferation

In order to further investigate the role of *hasp-1* in *C. elegans*, we generated a conditional degradation allele using the auxin-inducible degron (AID) system (Nishimura et al., 2009;Zhang et al., 2015). A previous study generated a *hasp-1* allele with a C-terminal AID tag to characterize *hasp-1* function in oocyte meiosis (Ferrandiz et al., 2018). We independently generated a similar allele, *hasp-1(pdx3)*, but with the AID tag and three FLAG tags on the N-terminus (hereafter referred to as *aid::hasp-1).* Mutant *aid::hasp-1* animals had no obvious phenotype in the absence of auxin, confirming that the tag does not alter protein activity or regulation. We crossed the *aid::hasp-1* allele into a background in which the auxin-activated ubiquitin ligase TIR1 is expressed in the germline under the control of the *sun-1* promoter (Zhang et al., 2015). Unlike the *hasp-1* deletion mutation (Fig. 1), HASP-1 protein degradation in the germline did not cause obvious changes germline morphology by DAPI staining (**Fig. 3A**), but did lead to an incompletely penetrant maternal-effect embryonic lethality (25% embryonic viability, **Fig. 3B** and **Table 1**). Maternal-effect embryonic lethality can be caused by defects in meiotic chromosome segregation and, consistent with this, the surviving embryos had a high incidence of males (Him) phenotype, which is caused by nondisjunction of the X chromosome in *C. elegans* (**Fig. 3B**). We monitored the development of embryos from mothers exposed to auxin and saw variable defects in the early cell divisions in embryos that did not hatch. Defects included early mitotic failures that generated abnormally large nuclei, cytokinesis failures or disorganized cleavage furrows producing multinucleate cells, and dead embryos arrested with various cell numbers well prior to gastrulation (**Fig. 3C**). Together with the Him phenotype, these data suggest that *hasp-1* is required both in meiosis and in the early mitotic divisions in embryos.

**Figure 3.**
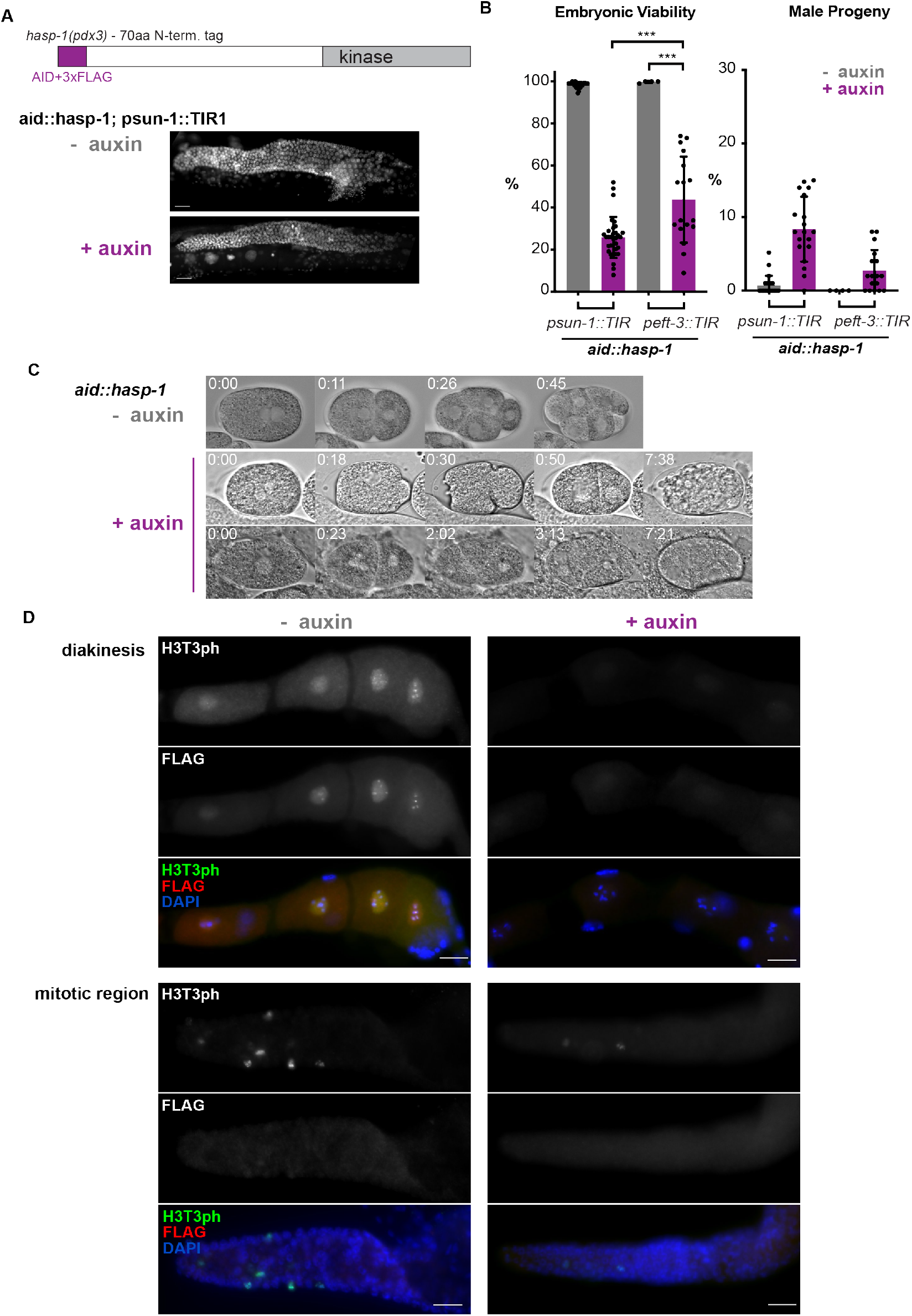
Analysis of HASP-1 degradation in the germline or soma. A) Schematic of the *aid::hasp-1* allele (top) and DAPI-stained germlines (below) in whole mount animals with or without exposure to auxin. B) Average embryonic viability and male production for whole broods (error bars show standard deviation) from *aid::hasp-1* animals in which the TIR1 ligase was expressed in the germline *(psun-1::TIR1)* or soma *(peft-3::TIR1).* *** = p<0.001 by Mann Whitney test. C) Nomarski images of embryos from *aid::hasp-1* animals with or without auxin treatment. Time shown is hr:min after pronuclear meeting. D) Immunofluorescence of dissected gonads from *aid::hasp-1* animals with or without auxin treatment. Scale bars = 10 μm.

**Table 1.** Brood analysis of *aid::hasp-1* animals.

| *genotype* | *auxin* | *Total % viability* | *% male* | *n (all eggs)* | *Ave. viability* | *Ave.% males* | *n (broods)* |
| --- | --- | --- | --- | --- | --- | --- | --- |
| *aid::hasp-1, p<sub>sun-1</sub>::TIR1* | - | 98.2 | 0.4 | 10120 | 98.1 | 0.4 | 30 |
| *aid::hasp-1, p<sub>sun-1</sub>::TIR1* | + | 25.2 | 8.6 | 6789 | 25.9 | 8.5 | 33 |
| *aid::hasp-1, p<sub>etf-3</sub>::TIR1* | - | 99.7 | 0* | 814 | 99.8 | 0 | 4 |
| *aid::hasp-1, p<sub>etf-3</sub>::TIR1* | + | 48.6 | 1.7 | 1883 | 43.7 | 2.7 | 17 |

\* No males were identified in this condition

Haspin proteins have been shown to play a few roles outside of mitosis and meiosis in other model systems, so we next used our *aid::hasp-1* allele to test whether *hasp-1* is needed outside of the germline. We considered the possibility that the gonad morphology defect seen in the *hasp-1* deletion mutation (Fig. 1) could be due to a requirement for *hasp-1* in somatic tissue during gonad development. We crossed the *aid::hasp-1* mutation into in a background in which TIR1 is expressed under the control of the *eft-3/eef1A. 1* promoter and *unc-54* 3’ UTR, which have been shown to drive expression in somatic tissues but not in the germline (Zhang et al., 2015). We saw no morphological defects after depleting *hasp-1* using this strain, but to our surprise, we observed a less penetrant maternal-effect embryonic lethality (44% embryonic viability) and a slight Him phenotype (**Fig. 3B** and Table 1). Because these phenotypes are similar to those observed upon germline knockdown, and are not expected to result from loss of HASP-1 in somatic tissue, we favor the conclusion that there are low levels of TIR1 protein in the germline that was not previously observed.

Next, we used immunofluorescence to better characterize *hasp-1* function in the germline. We could detect faint signal for the FLAG epitope in oocyte nuclei in the proximal region but not in mitotic nuclei in the distal gonad (**Fig. 3D**). The lack of detectable anti-FLAG signal in mitotic nuclei is surprising because the H3T3ph signal is robust and often covers all chromatin with high intensity, rather than being restricted to the mid-bivalent region as it is in oocytes. The H3T3ph and FLAG signals in oocytes were reduced below detectable levels when *aid::hasp-1* animals were exposed to auxin. There was also a clear reduction in H3T3ph staining in mitotic nuclei, however some residual H3T3ph could be detected in mitotic nuclei following auxin treatment. Thus, these data show a discrepancy between the level of HASP-1 protein detected by the anti-FLAG antibody and the level of H3T3ph, which may be evidence of a difference in HASP-1 activity in the two different cell types.

### Altering the timing of HASP-1 degradation separates phenotypes due to sperm and oocyte meiosis

Because the maternal-effect, dead-embryo phenotype we observed could be caused exclusively by the known role for *hasp-1* in oocyte meiosis, we next tested whether altering the time of auxin exposure could determine whether *hasp-1* was playing a role at earlier stages in the germline. All *aid::hasp-1* degradation experiments described above were done by transferring mothers to auxin during the L4 stage (when spermatogenesis is occurring), so we reasoned that earlier exposure to auxin could reveal roles in germline stem cell proliferation or spermatogenesis (**Fig. 4A**). When *aid::hasp-1* embryos were placed on plates containing auxin, they all hatched and grew into fertile adults (30/30), confirming that the embryonic lethality is due to knockdown in the mothers. When mothers were exposed to auxin as embryos, we found only a slight increase in the maternal-effect embryonic lethality relative to mothers exposed during the L4 stage (**Fig. 4B**). In contrast, exposing mothers to auxin later, after spermatogenesis but during oogenesis, caused similar levels of embryonic lethality as exposure starting during the L4 stage (**Fig. 4B**). Together, these data suggest that the most critical time of function of *hasp-1* is during adulthood, presumably during CPC recruitment in oocytes.

**Figure 4.**
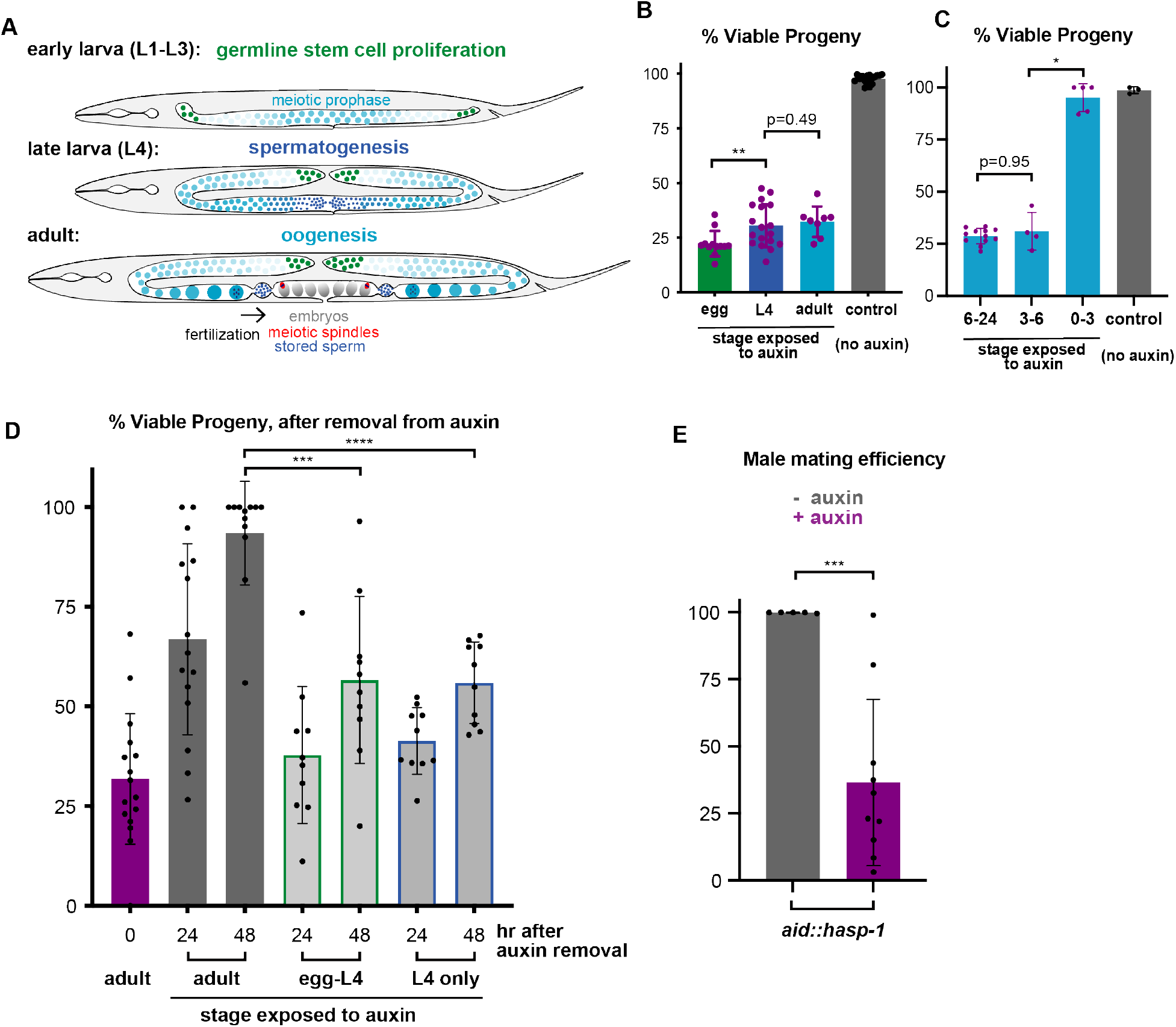
Time of function experiments show a requirement for *hasp-1* in spermatogenesis. A) schematic of worm showing stages of germline development relevant to auxin recovery experiments. B-D) Average embryonic viability of progeny lied by mothers treated with the time-course indicated. Each data point represents the viability for one plate with 5-20 worms, error bars show standard deviation. E) Mating efficiency of *aid::hasp-1* males crossed to control hermaphrodites. Each point is the number of cross progeny on a single mating plate. ****, ***, **, * = p < 0.0001, 0.001, 0.01, 0.05, respectively, by Mann Whitney test.

Pulse-labeling experiments have shown that oocytes likely progress from the diakinesis stage to embryos in the range of ~10 hr (Jaramillo-Lambert et al., 2007) so we tested whether a requirement for *hasp-1* could be seen following shorter exposures to auxin. Remarkably, embryos laid 3-6 hours after adult mothers were first exposed to auxin showed similar levels of lethality as embryos laid 6-24 hr after exposure **(Fig. 4C**). In contrast, very low lethality was seen in embryos laid in the first 3 hr after exposure. Previous work has shown that auxin-dependent degradation can occur in as little as 30 min using the same 1 mM auxin conditions used here and that most proteins can be degraded using this treatment for 4 hr (Divekar et al., 2021;Zhang et al., 2015). None of our degradation conditions were able to produce completely penetrant maternal-effect embryonic lethality, regardless of the length of treatment. The embryos that survive following maternal HASP-1 depletion presumably have achieved sufficient CPC recruitment, perhaps during prometaphase of meiosis I and dependent on another recruitment pathway.

Since we noticed that mothers exposed to auxin as embryos had lower embryonic viability than those exposed during the L4 stage or later, we reasoned that the difference was due to a role in sperm development, which occurs during the L4 stage in hermaphrodites (**Fig. 4A**). We performed auxin recovery experiments to test this hypothesis. We exposed hermaphrodites to auxin and then removed them and monitored whether the viability of their progeny recovered. When animals were exposed to auxin as adults (after spermatogenesis is over) for 24hr and then removed from auxin, their progeny recovered viability up to 93.5% after 48 hours (**Fig. 4D**). However, when animals were exposed to auxin from eggs and then removed at the end of the L4 stage, their progeny only recover to 56.7% after 48 hours. A similar lack of recovery (to 56.0% viability) was seen when animals were only exposed during spermatogenesis in the L4 stage. These results are consistent with a role for *hasp-1* in spermatogenesis because defective sperm would be stored and continue to cause dead embryos throughout an animal’s life, while oocyte meiosis is able to recover. To independently test whether *hasp-1* was required for functional sperm, we performed mating efficiency experiments in which *aid::hasp-1* males were mated to control hermaphrodites on plates with or without auxin. Consistent with a role for *hasp-1* in spermatogenesis, mating efficiency was severely reduced in the presence of auxin (**Fig. 4E**). While mating efficiency defects could be due to behavioral phenotypes, taken together our data are most likely explained by a role for *hasp-1* in CPC recruitment during spermatogenesis.

### Germline HASP-1 degradation shows genetic interactions with plk-2, bub-3 and sgo-1

In order to examine whether *hasp-1* interacts with known Haspin/CPC-regulators such as Pds5, Polo-like kinases, or the Bub1 pathway as characterized in other organisms, we tested genetic interactions between these mutants and HASP-1 degradation using the *aid::hasp-1* allele. First, we examined mitosis in germline stem cells by assaying germline proliferation. We reasoned that a genetic interaction with *hasp-1* could produce a germline proliferation defect similar to the *hasp-1* deletion strain (Fig. 1). We crossed *aid::hasp-1* into available mutant backgrounds that might impact HASP-1 activity and/or recruitment of the CPC (**Fig. 5A**), but found that none of the genetic backgrounds tested produced a synthetic germline proliferation defect. Previous work had shown that strong loss-of-function alleles of the Pds5 homolog *evl-14* cause germline proliferation defects and thus have small gonads in addition to defects in cohesion and vulval development (Wang, F. et al., 2003). We observed the small gonad phenotype in *evl-14* mutants as well as some disorganized nuclei, but this phenotype was not enhanced by HASP-1 degradation (**Fig. 5B**). Similarly, mutant backgrounds we tested that have incompletely penetrant maternal-effect embryonic lethality, *plk-2* and *bub-3*, did not have any obvious reduction in gonad size after HASP-1 degradation. As an alternative method to measure germline proliferation, we counted the total number of ovulations for individual animals as the sum of the number of eggs and unfertilized oocytes. Depletion using *aid::hasp-1* alone showed a significant reduction in ovulations but we saw no significant reductions when HASP-1 was depleted in mutant backgrounds relative to un-depleted controls (**Fig. 5C**). Thus, mitotic germline proliferation appears to be robust and less susceptible to failure in situations with low HASP-1 activity.

**Figure 5.**
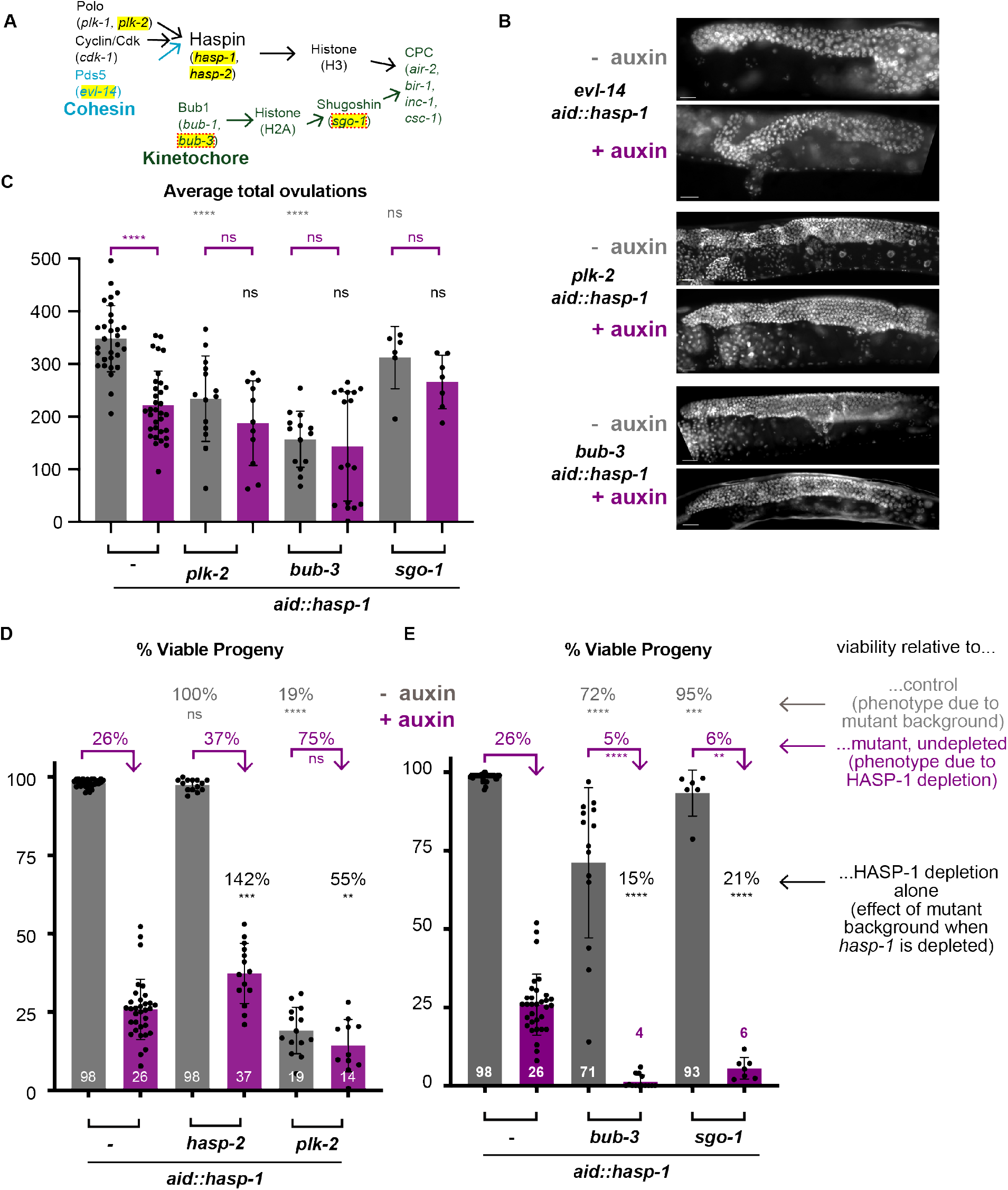
Degradation of *hasp-1* causes synthetic lethality with Bub pathway genes. A) Schematic of the Haspin and Bub1 pathways of CPC recruitment. Genes investigated in this figure are highlighted in yellow. B) Gonad nuclei in whole-mount DAPI-stained hermaphrodites of the indicated genotypes, with or without *hasp-1* depletion. Scale bars = 20 μm. C-E) Average total ovulations (sum of fertilized eggs and unfertilized oocytes), or % viability of whole broods. Statistical significance and percentages above bars show the relative viability compared to undepleted (-auxin) *aid::hasp-1* animals (grey), to undepleted *aid::hasp-1* in each mutant background (purple), or to *hasp-1* depleted (+ auxin) animals only (black). ns = not significant; ****, ***, **, * = p < 0.0001, 0.001, 0.01, 0.05, respectively, by Mann Whitney test.

Next, we analyzed maternal-effect embryonic lethality in the double mutant strains to determine if genetic interactions could be measured in embryonic viability. Embryonic lethality can result from failures in meiosis as well as mitosis during embryogenesis so this phenotype reports on the success of both mitosis and meiosis. The *hasp-2* or *plk-2* mutant backgrounds might further reduce the level of H3T3ph if *hasp-2* is able to substitute for *hasp-1* or if *plk-2* activates the HASP-1 protein. Consistent with the lack of phenotype in *hasp-2* mutants alone (Fig. 2), degrading HASP-1 in the *hasp-2* background did not enhance the embryonic lethality phenotype and instead caused an increase in embryonic viability (**Fig. 5D**). The *plk-2* mutant produces a strong embryonic lethality phenotype alone (Harper et al., 2011) but HASP-1 degradation did not cause a significant further enhancement of that phenotype (**Fig. 5D**). The *plk-2*, HASP-1-depleted animals had an embryonic viability that was greater than half (55%) of the viability caused by HASP-1 depletion alone, which was much less severe than the 19% viability that *plk-2* produced alone. An additive effect of the two mutations would have been expected to lower the viability in *plk-2*, HASP-1-depleted animals to below 5% (rather than the 14% we measured). This result suggests that *hasp-1* and *plk-2* are acting in the same pathway, which provides evidence that *plk-2* is able to activate the HASP-1 protein in *C. elegans.*

In contrast to our results with *plk-2*, we saw more severe genetic interactions when we performed HASP-1 degradation in backgrounds with mutations that should reduce Bub1-dependent recruitment of the CPC. The *C. elegans bub-1* locus is less than a map unit away from the *hasp-1* gene, so we crossed the *aid::hasp-1* mutation into a *bub-3* deletion mutant to indirectly disrupt the function of *bub-1.* Homozygous *bub-3(ok3437)* mutants have been shown to cause a strong reduction in the level of BUB-1 protein and to result in a slight embryonic lethality phenotype and a reduced brood size (Kim et al., 2015). In our experiments, the *bub-3* mutant had a mild embryonic viability defect (71%), which was very variable (SD = 24%) likely due in part to a large variability in brood sizes (**Table 2**). When *aid::hasp-1* was used in the *bub-3* background, this viability went down almost to zero, with a majority of broods having no viable eggs (11/17). Similarly, a deletion allele of the shugoshin homolog *sgo-1* (Ferrandiz et al., 2018) had only a slight embryonic viability phenotype alone (93%), but this dropped severely (to 6%) when HASP-1 was degraded. In both cases, the double mutant phenotype is well below what would be predicted for an additive effect. These data suggest that synergistic lethality is caused by the combined knockout of both the Haspin and Bub1 pathways of CPC recruitment, confirming that both pathways are important in *C. elegans.*

**Table 2:**
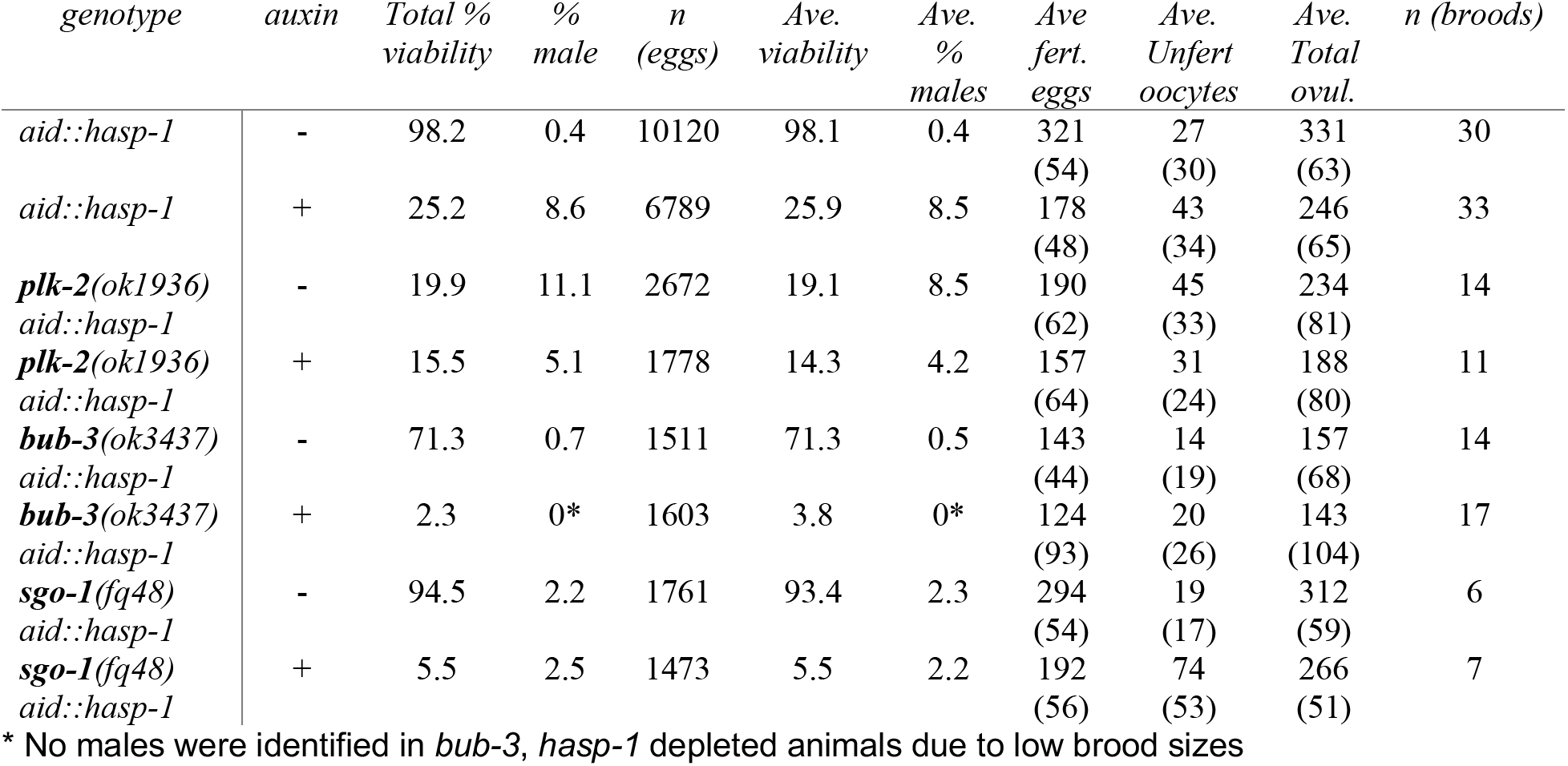
Brood analysis of double mutant strains.

## Discussion

### Ruling out a role for hasp-2 in C. elegans

Our genetic analysis provides evidence that the Haspin-related gene *hasp-1* acts during multiple processes in *C. elegans*, including oocyte meiosis, spermatogenesis, germline stem cell proliferation, and mitosis in early embryos (Fig. 1 & Fig. 3). We confirmed that *hasp-1* is the main ortholog of Haspin by ruling out roles for its most closely related paralog *hasp-2* (Fig. 2). The unusual expansion of the Haspin gene family in *C. elegans* (Higgins, 2001) has left open the possibility that sub functionalization has evolved among the different paralogs, but this does not appear to be the case for *hasp-2* in cell division. However, *hasp-2* and the other paralogs could retain the ability to phosphorylate histones, so further work will be needed to test whether these genes play important roles.

### Characterizing roles for hasp-1 during cell division in a variety of cell types

The adult sterility we observed in animals with a *hasp-1* deletion mutation is consistent with *hasp-1* being essential for cell division because in *C. elegans*, loss of function of cell division genes often produces sterility rather than embryonic lethality. Maternally deposited gene products are thought to be sufficient for cell division during embryonic development but then become limiting when the animals reach adulthood (O’Connell et al., 1998). Thus, the sterility phenotype of *hasp-1* deletion mutants does not provide evidence for other roles outside of cell division. However, our finding that *hasp-1* deletion mutations have severe defects in germline stem cell proliferation (Fig. 1) are in contrast with a recent report that found no defect in germline proliferation in Haspin knockout mice (Soupsana et al., 2021), showing that there are differences between animals in the importance of the Haspin pathway specifically in germline stem cells, as well as more generally in mitosis.

Because the *hasp-1* deletion mutation prevented formation of a normal germline, we used the AID system to further characterize roles for *hasp-1.* We were surprised to find that germline-specific depletion of HASP-1 did not recapitulate the severe germline stem cell proliferation defect of the *hasp-1* deletion, but instead produced animals with normal adult germline morphology (Fig. 3). HASP-1-depleted animals did produce fewer overall oocytes (Table 2), which could be due to a decrease in germline stem cell proliferation, but we did not see abnormal nuclei in the germline, a phenotype that has been produced by degradation of other essential cell division genes using the AID system (Zhang et al., 2015). Germline HASP-1 depletion did produce a variety of phenotypes consistent with roles for *hasp-1* in spermatogenesis, oocyte meiosis, and mitosis in embryos. The cell division defects we saw in embryos were similar to those seen following RNAi of CPC components (Kaitna et al., 2000;Romano et al., 2003;Speliotes et al., 2000) and the Him phenotype of the surviving progeny was consistent with a requirement for *hasp-1* in CPC localization in oocyte meiosis (Ferrandiz et al., 2018). The low embryonic viability we measured is therefore due to a combination of meiotic and mitotic defects.

It is notable that the germline knockdown of HASP-1 produced more severe phenotypes in oocytes than during germline mitotic divisions. This may support the possibility that the relative importance of *hasp-1* differs in these two cell types. HASP-1’s contribution to CPC recruitment may be dispensable during the mitotic divisions even though it is required in spermatogenesis (Figure 4) and in oocyte meiosis (Ferrandiz et al., 2018;Rogers et al., 2002). CPC recruitment by HASP-1-independent mechanisms, such as the activity of Bub1, may be able to more fully compensate for loss of *hasp-1* in germline stem cell mitosis than in meiosis or mitosis in embryos. Alternatively, the different severity of HASP-1 degradation phenotypes could simply be because the level of HASP-1 activity was reduced more completely in meiosis than in mitosis, which is supported by the residual levels of H3T3ph we were able to detect after HASP-1 depletion (Fig. 3). However, even if this is true, our data still show a potentially interesting difference in HASP-1 activity in germline stem cells relative to oocytes because it appeared that undetectable levels of HASP-1 protein in mitotic nuclei were able to generate higher H3T3ph (Fig. 3). The apparent decrease in HASP-1 activity may be a result of mechanisms that restrict HASP-1 protein to a small region of the chromosome in meiosis to support differential cohesion release, or to higher levels of phosphatase activity on meiotic chromosomes (de Carvalho et al., 2008;Kaitna et al., 2002;Rogers et al., 2002;Tzur et al., 2012).

Further work will be needed to more precisely quantify the levels of HASP-1 protein, H3T3ph, and CPC recruitment. Our data suggest that HASP-1 protein levels are very low in the *C. elegans* germline (Fig. 3), consistent with what has been observed for Haspin in other organisms with the exception of mouse spermatocytes (Tanaka et al., 1999). One explanation for the surprising meiotic defects we observed following HASP-1 degradation in somatic tissue (Fig. 3) is that the low level of HASP-1 protein makes knockdown is possible with low levels of TIR1 protein in the germline. Quantification of H3T3ph was complicated by technical challenges with H3T3ph antibodies. In our hands, H3T3ph antibodies must be used at low concentrations to avoid staining throughout the nucleoplasm, which we found to be present with a variety of fixation conditions (data not shown). Nucleoplasmic staining did not appear to be an aftifact because it was specific to the proximal oocytes where chromosomal H3T3ph is expected, and was reduced when animals were exposed to auxin. The nucleoplasmic signal therefore may reflect phosphorylation of the soluble histone pool, since soluble HASP-1 is unlikely to be inhibited by the recruitment of phosphatases by LAB-1, which is localized to the chromosome axis (de Carvalho et al., 2008; Ferrandiz et al., 2018).

### Genetic interactions within and between the Haspin and Bub1 pathways

Combining AID-based degradation of HASP-1 with other mutant backgrounds allowed us to support models for the activation of HASP-1 protein and the recruitment of the CPC. The parallel roles of Haspin and Bub1 pathways in CPC recruitment were supported by genetic interactions in *S. pombe* and we set out to take a similar approach (Yamagishi et al., 2010). We found that HASP-1 depletion in the *plk-2* background caused a less severe phenotype than would have been expected from the additive phenotypes caused by the loss of function of each alone, which is consistent with *hasp-1* and *plk-2* acting in the same pathway. This conclusion is also based on the evidence from Xenopus and human cells that Plk1 phosphorylates Haspin to activate it (Ghenoiu et al., 2013; Zhou et al., 2014). Our results provide the most direct evidence to date that *plk-2* is able to activate the HASP-1 protein in *C. elegans* and that this regulatory mechanism is conserved in this model system.

In contrast to our results with *hasp-1* and *plk-2*, we saw synthetic interactions between *hasp-1* and the Bub1 pathway components *bub-3* and *sgo-1.* These results were somewhat surprising because previous work in *C. elegans* showed that depletion of *hasp-1* using a similar degron allele completely removed CPC localization in diakinesis oocytes and that *sgo-1* deletion did not affect CPC recruitment (Ferrandiz et al., 2018). However, although the Bub1 pathway is therefore not active in recruitment of the CPC in diakinesis, our data provide evidence that there is still a role for the Bub1 pathway in CPC recruitment in oocytes, perhaps later in meiosis after BUB-1 protein is recruited to kinetochores and the ring that forms at the mid-bivalent (Dumont et al., 2010). Previous work on the function of meiotic axis proteins has shown that the CPC can still be rapidly recruited to chromosomes in prometaphase in mutants that fail to recruit the CPC in diakinesis (Sato-Carlton et al., 2018). Thus, further work on CPC recruitment in *C. elegans* will help tease out the robust mechanisms that recruit this important complex.

## Materials and Methods

### Worm strains and maintenance

All worms were maintained on MyoB plates seeded with OP50 bacteria. All crosses were conducted at 20 °C and genotypes were monitored by PCR. The deletion allele of *hasp-1* was obtained from the Japanese Bioresource Center (Gengyo-Ando and Mitani, 2000). Worm strains used in this study were as follows.

DJW05 – *hasp-2(pdx1)* I.

DJW31 - *hasp-1(pdx3* [3xflag::aid::*hasp-1*]) I; *ieSi38 [sun-1p::TIR1::mRuby::sun-1 3’UTR + Cbr-unc-119(+)]* IV.

DJW32 - *hasp-1(pdx3 [3xflag::aid::hasp-1])* I; *ieSi57 [eft-3p::TIR1::mRuby::unc-54* 3’UTR + Cbr-*unc-119(+)]* II.

DJW36 - *hasp-2(pdx1) hasp-1(pdx3 [3xflag::aid::hasp-1])* I; *ieSi38 [sun-1p::TIR1::mRuby::sun-1* 3’UTR + Cbr-*unc-119*(+)] IV.

DJW39 - *hasp-1*(pdx3) I; *bub-3(ok3437)* II; *ieSi38 [sun-1p::TIR1::mRuby::sun-1* 3’UTR + Cbr-*unc-119(+)]* IV.

DJW42 - *plk-2(ok1936) hasp-1(pdx3)* I; *ieSi38 [sun-1p::TIR1::mRuby::sun-1* 3’UTR + Cbr-*unc119(+)]* IV.

DJW47 - *hasp-1*(pdx3) I; *ieSi38 [sun-1p::TIR1::mRuby::sun-1* 3’UTR + Cbr-*unc119*(+)] *sgo-1(fq48)* IV.

DJW49 - *hasp-1*(pdx3) I; *unc-36(e251) evl-14(ar112)/dpy-17(e164) unc-32(e189)* III. *ieSi38 [sun-1p::TIR1::mRuby::sun-1* 3’UTR + *Cbr-unc-119(+)]* IV.

DJW64 - *hasp-1* (pdx3)/*tmC18 [dpy-5(tmIs1236)]* I. *ieSi38 [sun-1p::TIR1::mRuby::sun-1 3’UTR + Cbr-unc-119(+)] ltIs37 [pAA64; pie-1/mCherry::his-58; unc-119(+)]* IV. *qIs56 [lag-2p::GFP + unc-119(+)]* V.

DR429 - *dpy-5(e61) unc-15(e73)* I.

N2 – wild type

### Genome editing

New alleles were generated by injection of Cas9-gRNA RNP complexes using *dpy-10* Co-CRISPR to enrich for edited progeny (Arribere et al., 2014;Paix et al., 2015). Cas9-NLS protein was complexed *in vitro* with equimolar quantities of duplexed tracrRNA and target-specific crRNAs. All RNP components were purchased from IDT. Repair templates were single-stranded Ultramer or Megamer oligonucleotides purchased from IDT. Each 10 μl of injection mixture contained: 16 mM [Cas9 protein + gRNA] (target gene + *dpy-10*), plus 0.5–1.5 μM repair templates. Injected animals were transferred to individual plates, incubated at 20 °C, and screened for F1 Rol or Dpy progeny.

Disruption of the *hasp-2* gene was accomplished by inserting the sequence GCTAG after the first base in the 14^th^ codon, introducing a stop codon, a frameshift, and an Nhe I restriction site. The *aid::3xFLAG::hasp-1* allele was generated by inserting a 70aa tag (based on those used in (Zhang et al., 2018)) after the 3^rd^ codon of *hasp-1*, full sequence shown below.

3xFLAG::AID tag (with GAGS spacer in between the FLAG and AID, lower case) GATTATAAAGACCATGATGGAGACTATAAGGATCACGATATTGATTACAAAGACGATGATGA TAAAggagccggatctCCTAAAGATCCAGCCAAACCTCCGGCCAAGGCACAAGTTGTGGGATGG CCACCGGTGAGATCATACCGGAAGAACGTGATGGTTTCCTGCCAAAAATCAAGCGGTGGC CCGGAGGCGGCGGCGTTCGTGAAG

### Protein sequence analysis

Wild-type *C. elegans* gene sequences were obtained in FASTA format from Wormbase (www.wormbase.org). For the HASP-1 and HASP-2 protein sequences, the conceptual translations from Wormbase were used, while homologous human, mouse, *S. pombe*, and S. cerevisiae sequences were obtained from the NCBI database (https://www.ncbi.nlm.nih.gov/protein/). Accession numbers are as follows: human Haspin (NP_114171.2), mouse (BAB00640.1), *S. pombe* (CAB16874.1), *D. melanogaster* (NP_001015349.2). All alignments were done using Clustal Omega (Sievers et al., 2011) and visualized using Jalview (Waterhouse et al., 2009).

### Embryonic viability, male frequency, and brood size assays

Percent viability, male frequency, and brood size of self progeny were measured by placing single L4 hermaphrodites onto 60 mm plates at 20°C and transferring worms to a new plate every 24 h until they stop laying. Dead eggs were scored 24 h after the mother had been removed and the number and sex of F1 were scored after they reached the adult or L4 stages. The embryonic viability for each worm was calculated by dividing the total number of progeny by the total number of worms and dead eggs laid while percent males was the number of males divided by the total progeny. In all graphs, the % viability shown is the average for all worms for which the entire brood was counted. Brood size was the total number of worms and dead embryos, and the total ovulations was the sum of worms, eggs, and unfertilized oocytes.

### Male mating assays

Mating efficiency was measured by placing six dpy-5(e61) unc-15(e73) L4 hermaphrodites on 60 mm plates with six males. Males and hermaphrodites were allowed to mate for 48 h at 20°C. Males were removed from the plate, and hermaphrodites were transferred to new plates. Hermaphrodites were then transferred to new plates every 24 h until egg laying stopped. Plates were scored at 72 h of age. The number of wild type (outcross) and Dpy Unc (self) offspring were counted for each plate, and male mating efficiency was calculated by dividing the number of outcross offspring by the total number of offspring.

### Immunofluorescence

For immunostaining, dissected gonads were fixed in 2% formaldehyde in egg buffer containing 0.1% Tween 20 for 5 min, freeze cracked into cold methanol, and transferred to PBS + 0.1% Tween 20 at room temperature. Primary antibodies used were anti-phospho-Histone H3 (Thr3) (EMD Millipore 07-424) diluted 1:20,000 and anti-HTP-3 (gift from A. F. Dernburg, (MacQueen et al., 2005)) diluted 1:500. Secondary antibodies were purchased from Jackson ImmunoResearch Laboratories, Inc. DAPI staining was done using 0.5μg/mL in PBS on both dissected gonads or whole mount animals after cold methanol fixation. Samples were mounted in Prolong Gold mounting medium (Millipore Sigma).

### Microscopy

Confocal imaging was performed on a Diskovery spinning disk confocal system (Andor) mounted on a Nikon Eclipse Ti microscope with a 60× 1.4 NA objective. Epifluorescence and DIC (Nomarski) imaging were performed on an Olympus BX60 microscope with a Plan Apo 60x 1.42 NA objective.

## Acknowledgements

We would like to thank N. Peel for use of the confocal microscope; A. F. Dernburg for the HTP-3 antibody; A. Sato-Carlton and E. Martinez-Perez for helpful comments on the manuscript; and N. Shao and E. Martinez-Perez for sharing unpublished observations that corroborated and enhanced our understanding of our results. Some strains were provided by the Caenorhabditis Genetics Center (CGC), which is funded by NIH Office of Research Infrastructure Programs (P40 OD010440). The *hasp-1* deletion strain was provided by the National BioResource Project (Japan).

## Author contributions

D.J.W. conceived and designed the experiments. J.M., I.R., M.W., D.C., M.S., A.M., E.H., K.B., and D.J.W. performed the experiments and analyzed the data. D.J.W. and J. M. wrote the paper.

